# Leptin and adiponectin dynamics at patients with rectal neoplasm - gender differences

**DOI:** 10.1101/541557

**Authors:** Alexandru Florescu, Dumitru Branisteanu, Stefana Bilha, Dragos Scripcariu, Ioana Florescu, Viorel Scripcariu, Gabriel Dimofte, Ioana Grigoras

## Abstract

**Background.:** Numerous studies associate adipokines with colorectal malignancy, but few data deal with patients suffering exclusively of rectal carcinoma (RC). Aims. We evaluated leptin and adiponectin levels in RC patients compared to healthy population and their dynamics after surgery. Material and methods. Serum leptin and adiponectin were evaluated before surgery in 59 RC consecutive patients (38 males and 21 females), and in age and weight matched healthy controls. Measurements were repeated at 24, 72 hours and 7 days after surgery. Results. Adipokine levels were higher in women. Controls had higher leptin (32.±4.34 vs 9.51±1.73 ng/ml in women and 11±2.66 vs 2.54±0.39 ng/ml in men, p=0.00048 and 0.0032) and lower adiponectin (9±0.64 vs 11.85±1.02 µg/ml in women and 7.39±0.51 vs 8.5±0.62 µg/ml in men, p=0.017 and 0.019) than RC patients. Surgery caused an increase of leptin from 5.11±0.8 to 18.7±2.42 ng/ml, p=6.85 × 10^8^, and a decrease of adiponectin from 9.71±0.58 to 7.87±0.47 µg/ml, p=1.4 × 10^10^ for all RC patients and returned thereafter to the initial range at 7 days. Adipokines were correlated with body weight (BW). The significance of correlation persisted after surgery only in males, but disappeared in females. Adipokines were not modified by tumor position, presurgical chemoradiotherapy or surgical technique. Women with RC experiencing weight loss had higher adiponectin than women without weight modifications (p<0.05 at all time points). Conclusions. Adipokine levels of patients with RC differ from the healthy population, possibly reflecting an adaptation to disease. Adipokine modifications after surgery may be related to acute surgical stress. Whether leptin and adiponectin directly interact is not clear. Women have higher adipokine levels, more so after significant weight loss, but the strength of their correlation with BW decreases after surgery. These data suggest gender differences in the adipokine profile of RC patients which may find clinical applications.

## Introduction

Adipose tissue performs an endocrine role, mediated by hormones (adipokines) which are specifically secreted by the adipocytes [1]. Epidemiological data consistently show an association between weight gain and the risk of colorectal cancer in adults [2,3]. This observation led to the evaluation of a possible linking role of adipokines with cancer pathogenesis [1-3]. Leptin is an adipokine which reflects fat mass, being involved in nutrition behavior, and is considered a proinflammatory and carcinogenic factor in various cancer types, stimulating carcinogenic angiogenesis [1,3-5]. Adiponectin is inversely related to fat mass and stimulates insulin secretion [1] and has an anti-inflammatory role, inhibiting angiogenesis and being considered protective against various cancer types [1,3-5]. Therefore adipokines are interesting candidates as biomarkers in colorectal cancer (CRC) with potential significance in cancer development and prognostic evaluation.

Literature shows conflicting data regarding leptin and adiponectin levels in digestive cancers. Different authors describe increased [1,2,6-8], decreased [9-13] or unchanged leptin levels [3,14], and – similarly – increased [15], decreased [1-3,8,16,17] or unchanged adiponectin levels [18,19] in patients with CRC. These discrepancies may be due to the fact that carcinogenesis is not only related to serum adipokine concentrations but also to adipokine receptors expressed on various cells, i.e. tissue sensitivity [3,5,11,19,20]. It is therefore unclear whether modifications in adipokine levels described in patients with digestive cancer represent a causative link or are merely an adaptive epiphenomenon [1,3,5,11,14,20]. Finally, the contrasting data described in the literature may be caused by the heterogeneity of studied groups with respect to tumor localization [2,8,11,12,17,19-22], gender [6,7,12,23], weight modification [2,9], disease severity [8,11,14], chemotherapy [4] or due to statistical complexity caused by multicentricity of the study or complex metaanalysis [2,3,9,17,19-23]. In order to avoid these variations, one needs to choose a more homogenous series, in a single location at similar stages. Such type of neoplasia is the carcinoma located in the rectal region (rectal carcinoma, RC) which is usually not accompanied by cachexia, a problem that might affect changes in adipokines.

The aims of our study were therefore to observe differences in leptin and adiponectin levels between patients suffering of RC operated in the same surgical center and healthy age, weight and gender-matched controls. We also intended to follow the evolution of leptin and adiponectin after surgery and at distance, at different time points.

## Materials and Methods

### Study participants

The study was approved by the Ethical Committee of Clinical Research of the Regional institute of Oncology of Iasi (nr. 167 from 30/06/2017). All study participants signed a written informed consent.

We recruited an initial number of 62 patients (40 males and 22 females) diagnosed with stage 2 and 3 RC and consecutively programmed for surgical intervention at the Regional Institute of Oncology of Iasi. Thirty patients followed neoadjuvant chemoradiotherapy (nCRT) for 2 months before surgery (pelvic irradiation of 45 Gy and concomitant Capecitabine 1600 – 1650 mg/m2/day) [24] and were operated 60 ± 10 days after chemoradiotherapy arrest, but two males and one female of chemoradiotreated patients dropped out from the study before surgery. The other 32 patients were operated directly, without nCRT. Patients were submitted to various techniques of surgical intervention, according to location within the rectum: low anterior resection type operations (LAR) in 42 patients and abdomino-perineal resection type (APR) in other 17 patients. Blood samples were collected before surgery, as well as at 24 hours, 72 hours and 7 days after surgery, in order to evaluate serum leptin and adiponectin levels. Other 20 patients (10 males and 10 females) gradually dropped out during the follow up period, due to refusal to further participate, withdrawal from the informed consent or absence at routine check-up 7 days after surgery (Fig 1).

**Fig 1.**
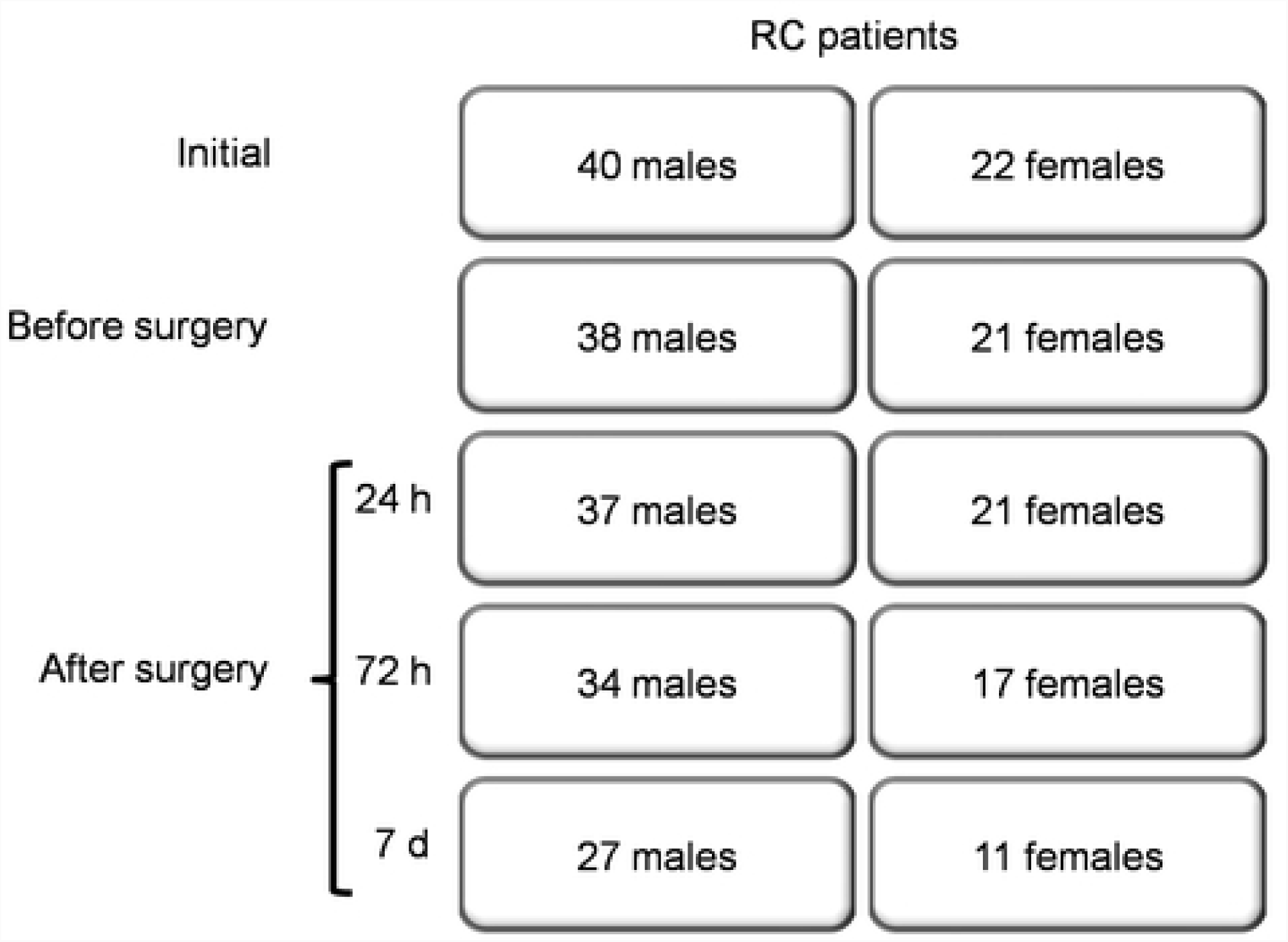
Recruitment and dropout diagram. RC = rectal carcinoma.

Serum leptin and adiponectin were also evaluated in 29 male and 28 female age and weight-matched healthy controls, with no malignant antecedents. All patients and controls were weighed with the same weighing scale. Patients with RC were questioned about their body weight (BW) evolution during the year before enrollment. The characteristics of RC patients and of controls are depicted in Table 1.

**Table 1.** Characteristics of RC patients and controls.

|  | Males |  |  | Females |  |  |
| --- | --- | --- | --- | --- | --- | --- |
|  | RC | Control | p | RC | Control | p |
| Number | 38 | 29 | - | 21 | 28 | - |
| Age<br>(years, mean ± SD) | 66.1 ± 8.3 | 63.9 ± 6.8 | 0.203 | 67.9 ± 8.4 | 66.8 ± 5.5 | 0.469 |
| Weight<br>(kg, mean ± SD) | 77.9 ± 11.3 | 81.5 ± 19.7 | 0.273 | 72.2 ± 11.4 | 76.5 ± 14.7 | 0.221 |

### Evaluation of serum leptin and adiponectin

After overnight fasting, a sample of 10 ml blood was drawn from forearm vein from all study participants and the separated serum aliquots were stored at −20°C until assessment. Blood collection of RC patients was performed before surgery and repeated 24 hours, 72 hours and seven days after surgery. Serum assessment of leptin and adiponectin was performed by using Luminex technology (Luminex Screening Assay, R&D Systems Inc., MN, USA).

### Statistical analysis

Data are expressed as mean ± standard error of the mean (SEM). SPSS (statistics version 20.0 for Windows) was used for statistical analysis. Shapiro-Wilk test was used to verify the normal distribution of data. Differences between groups were tested using the student’s t test and the Mann-Whitney U test. Correlation analysis (Pearson analysis for normally distributed data and Spearman rank correlation for skewed data) was performed to investigate the relationship between leptin and adiponectin, as well as between BW and leptin or adiponectin, respectively. The possibility of BW of being a confounder in the relationship between leptin and adiponectin was investigated through hierarchical regression. Differences were considered significant at p values < 0.05.

## Results

### Comparison of mean leptin and adiponectin levels between RC patients and healthy controls

Females had higher leptin and adiponectin levels than males in both control and RC groups. When compared to controls, RC patients had significantly lower leptin and higher adiponectin levels in both sexes (Table 2).

**Table 2.** Differences in adipokine values between patients with RC and controls and between males and females

| Parameter | CTR males | RC males | p values<br>CTR vs RC<br>males | CTR females | RC females | p values<br>CTR vs RC<br>females |
| --- | --- | --- | --- | --- | --- | --- |
| Leptin<br>(ng/ml) | 11 ± 2.6 *** | 2.5 ± 0.4** | ***p=0.0032*** | 32.2 ± 4.3 | 9.5 ± 1.7 | ***p=0.00049*** |
| Adiponectin<br>(µg/ml) | 7.4 ± 0.5* | 8.5 ± 0.6** | ***p=0.019*** | 9 ± 0.6 | 11.9 ± 1 | ***p=0.0171*** |

CTR – healthy control groups; RC – patients with rectal carcinoma. \* $p < 0.05$ , \*\* $p < 0.01$ , \*\*\* $p < 0.001$ compared to females.

### Dynamics of leptin and adiponectin after surgical intervention for RC

Leptin and adiponectin of RC patients showed a mirror profile after surgery. They displayed a steep increase of leptin (from 5.11 ± 0.8 to 18.7 ± 2.42 ng/ml, p = 6.85 × 10-8) and a significant decrease of adiponectin (from 9.71 ± 0.58 to 7.87 ± 0.47 μg/ml, p = 1.4 × 10-10) 24 hours after surgery. The levels of the two adipokines recovered to presurgical levels during the follow up period. The dynamics of the two adipokines is preserved when data are analyzed separately in the male and female subgroups (Fig. 2).

**Fig 2.**
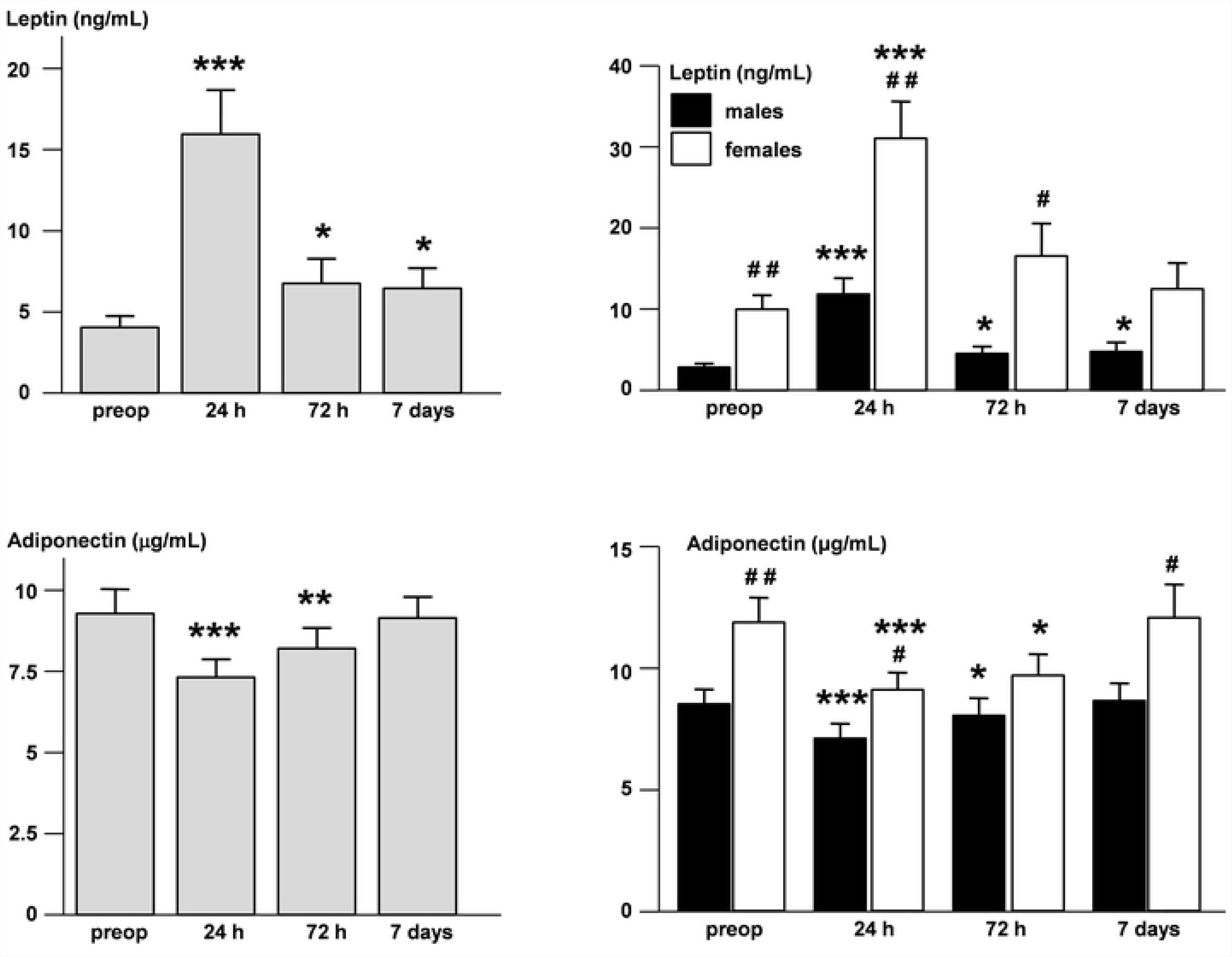
Dynamics of leptin and adiponectin after surgery. Leptin (upper panel) and adiponectin (lower panel) levels (mean +/-SEM) in all patients with RC (left, grey bars) and in males (right, black bars) or females only (right, white bars) before surgery and at 24, 72 hours and 7 days after surgery. * compared to presurgical values. # compared to males, *** p < 0.0001, ** or ## p < 0.01, * or # p < 0.05.

### Correlations between leptin, adiponectin and BW

BW did not change significantly during the follow-up week. Adipokines were correlated to BW before surgery in both males and females. Leptin was positively correlated, whereas adiponectin was inversely correlated. Surgery caused a persistent loss of correlation significance of leptin and adiponectin with BW in females, but not in males (Table 3).

**Table 3.** Correlation coefficients of BW with leptin and adiponectin in males and females with RC

| Gender | time | Leptin |  | Adiponectin |  |
| --- | --- | --- | --- | --- | --- |
| | | $r^2$ | p | $r^2$ | p |
| Males | Before surgery | 0.13 | ***0.027*** | 0.11 | ***0.045*** |
|  | 24 h | 0.132 | ***0.031*** | 0.159 | ***0.016*** |
|  | 72 h | 0.139 | ***0.028*** | 0.116 | ***0.048*** |
|  | 7 days | 0.322 | ***0.00056*** | 0.144 | ***0.049*** |
| Females | Before surgery | 0.263 | ***0.021*** | 0.239 | ***0.025*** |
|  | 24 h | 0.045 | 0.359 | 0.132 | 0.106 |
|  | 72 h | 0.088 | 0.247 | 0.15 | 0.125 |
|  | 7 days | 0.041 | 0.549 | 0.135 | 0.267 |

$r^2$ = regression coefficient of determination. With italic bold – significant correlations.

The gender difference of correlation slopes was obvious for the leptin-BW correlation at all post-surgical time points (Fig 3).

**Fig 3.**
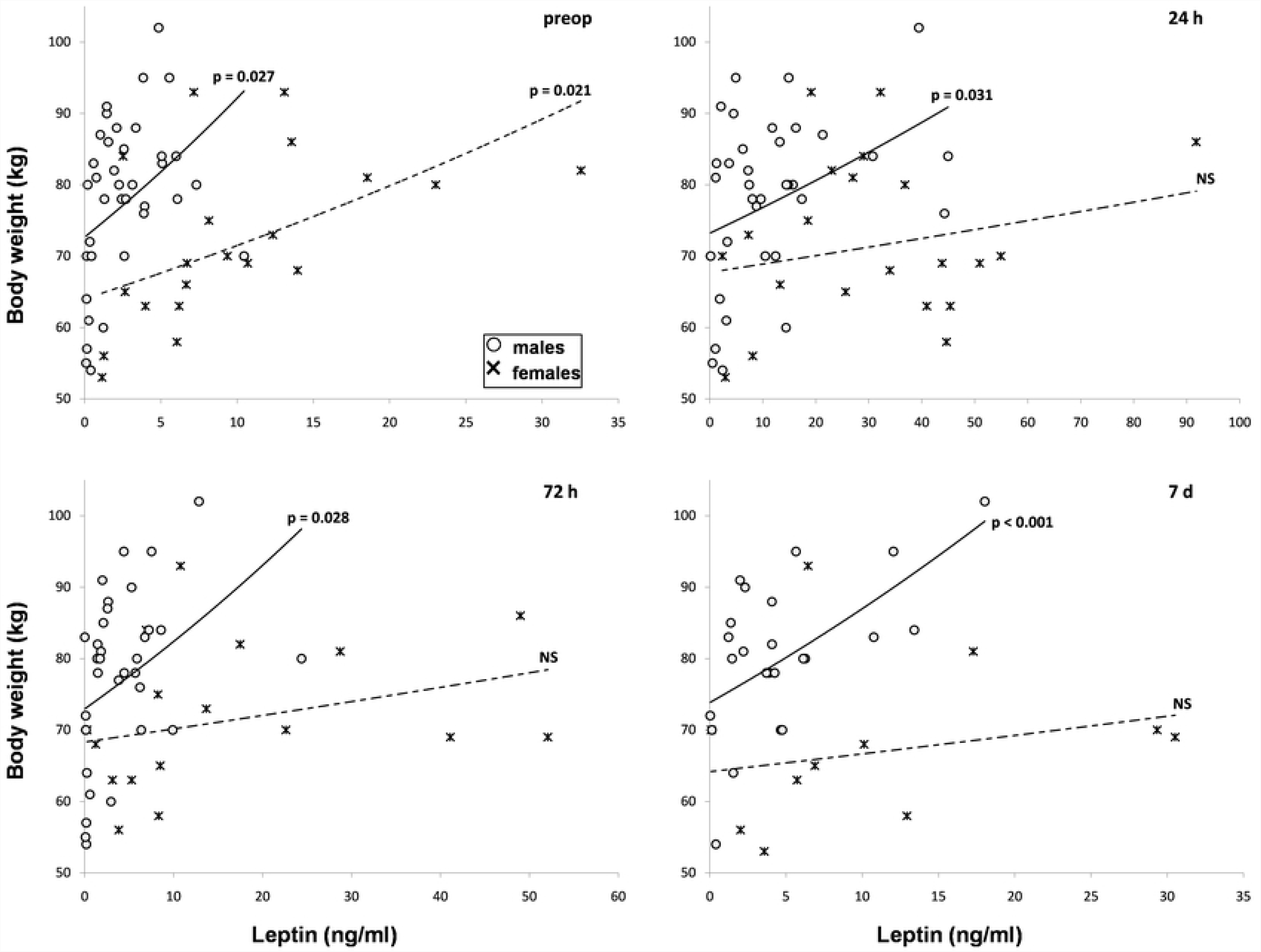
Correlations between weight and serum leptin in males (○) and females (×). Correlations before surgery (preop, up, left), after 24 hours (up, right), 72 hours (low, left) and seven days (low, right). Correlation was considered significant at p values lower than 0.05. NS = non-significant.

### Correlations between leptin and adiponectin

Leptin and adiponectin levels before surgery were inversely correlated in both male and female RC patients. Correlative significance was lost 24 hours after surgery, but was later regained one week after surgery in males and as soon as 72 hours after surgery in females (Fig 4).

**Fig 4.**
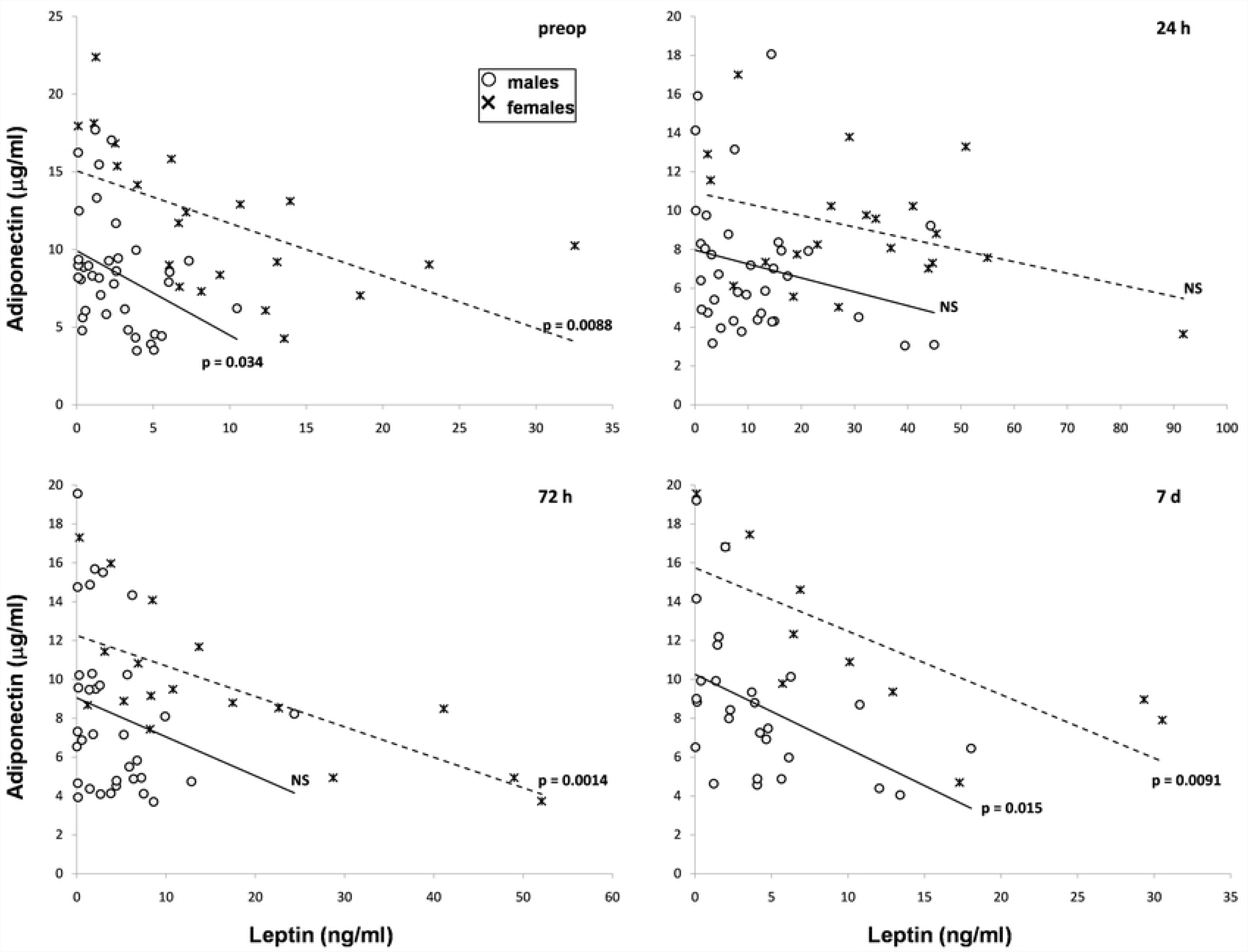
Correlations between serum leptin and adiponectin in males (○) and females (×) with CR cancer. Upper left – before surgery, upper right – 24 hours, lower left – 72 hours and lower right – one week after surgery. Correlation was considered significant at p < 0.05. NS = non-significant.

### Hierarchical regression analysis

The correlation of leptin and adiponectin with BW, as well as between leptin and adiponectin were significant before surgery in both males and females, and again at one week after surgery only in males, when neither leptin nor adiponectin were correlated with BW in females. In order to check whether the associations between leptin and adiponectin are independent or confounded by the BW, hierarhical regression analysis was performed (Table 4).

**Table 4.** Results of hierarchical regression.

| Gender | Dependent variable | Model | Predictor variable | $\beta$ | SEE | p |
| --- | --- | --- | --- | --- | --- | --- |
| M | adiponectin preop | 1 | leptin | **-0.351** | 0.865 | **0.034** |
|  |  | 2 | leptin | -0.265 | 0.260 | 0.123 |
|  |  |  | weight | -0.235 | 0.055 | 0.17 |
|  | adiponectin 7 d | 1 | leptin | **-0.468** | 0.145 | **0.014** |
|  |  | 2 | leptin | -0.378 | 0.187 | 0.111 |
|  |  |  | BW | -0.144 | 0.071 | 0.533 |
| F | adiponectin preop | 1 | leptin | **-0.57** | 0.112 | **0.007** |
|  |  | 2 | leptin | **-0.571** | 0.121 | **0.012** |
|  |  |  | BW | 0.005 | 0.053 | 0.981 |

M = males, F = females, preop = preoperative, BW = body weight, $\beta$ = standardized beta coefficient, SEE = standard error of the estimate, p = significance

Leptin (in the first step) and BW (added to the model in the second step) were entered as predictor variables for adiponectin levels. Standardized β coefficient for leptin decreased and became non-significant when BW was added to the model only in men before and at 7 days after surgery, but not in women before surgery, where the association between adiponectin and leptin remained significant after BW was added in the second step (p = 0.012).

### Influence of tumor localization, nCRT, type of surgery and weight loss on adipokine profile

The rectal tumor was located either in the lower rectum in 30 patients (21 males and 9 females) and in the upper rectum in the other 29 patients (17 males and 12 females). Thirty-two of enrolled RC patients (20 males and 12 females) did not receive nCRT before surgery, whereas 27 patients (18 males and 9 females) received chemoradiation using the long course protocol and operated 60 days ± 10 days after nCRT arrest. Forty-two patients (25 males and 17 females) were operated using LAR type surgery, whereas 17 patients (13 males and 4 females) were submitted to APR. Twenty-five patients (17 males and 8 females) experienced significant weight loss (between 3 and 30 kg in the last year, mean weight loss of 9.5 ± 1.4 kg for all 25 patients, and of 9.9 ± 1.5 and 8.8 ± 3.2 kg for males and females, respectively). The other 34 patients (21 males and 13 females) did not lose weight during the last year before surgery. Leptin and adiponectin had similar dynamics in all subgroups irrespective of tumor localization, the use of nCRT, the surgical technique used or the presence or absence of weight loss. (Table 5).

**Table 5.**
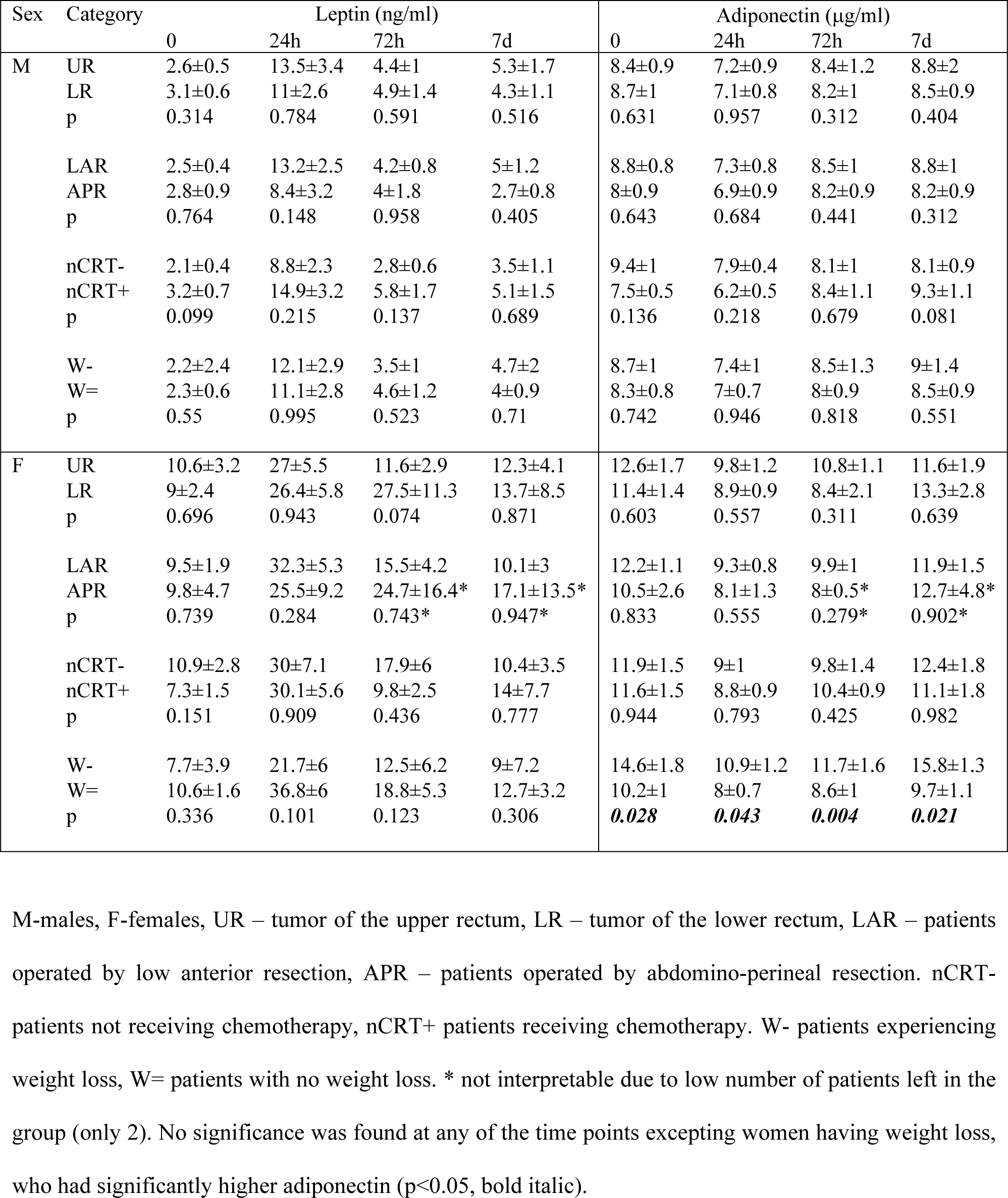
Adipokine levels at different tumor localizations, different surgery procedures, and in the presence or absence of presurgical nCRT or weight loss.

There were no significant differences between adipokine levels at different time points in both sexes, except for adiponectin, which had significantly higher levels at women who lost weight (Fig 5).

**Fig 5.**
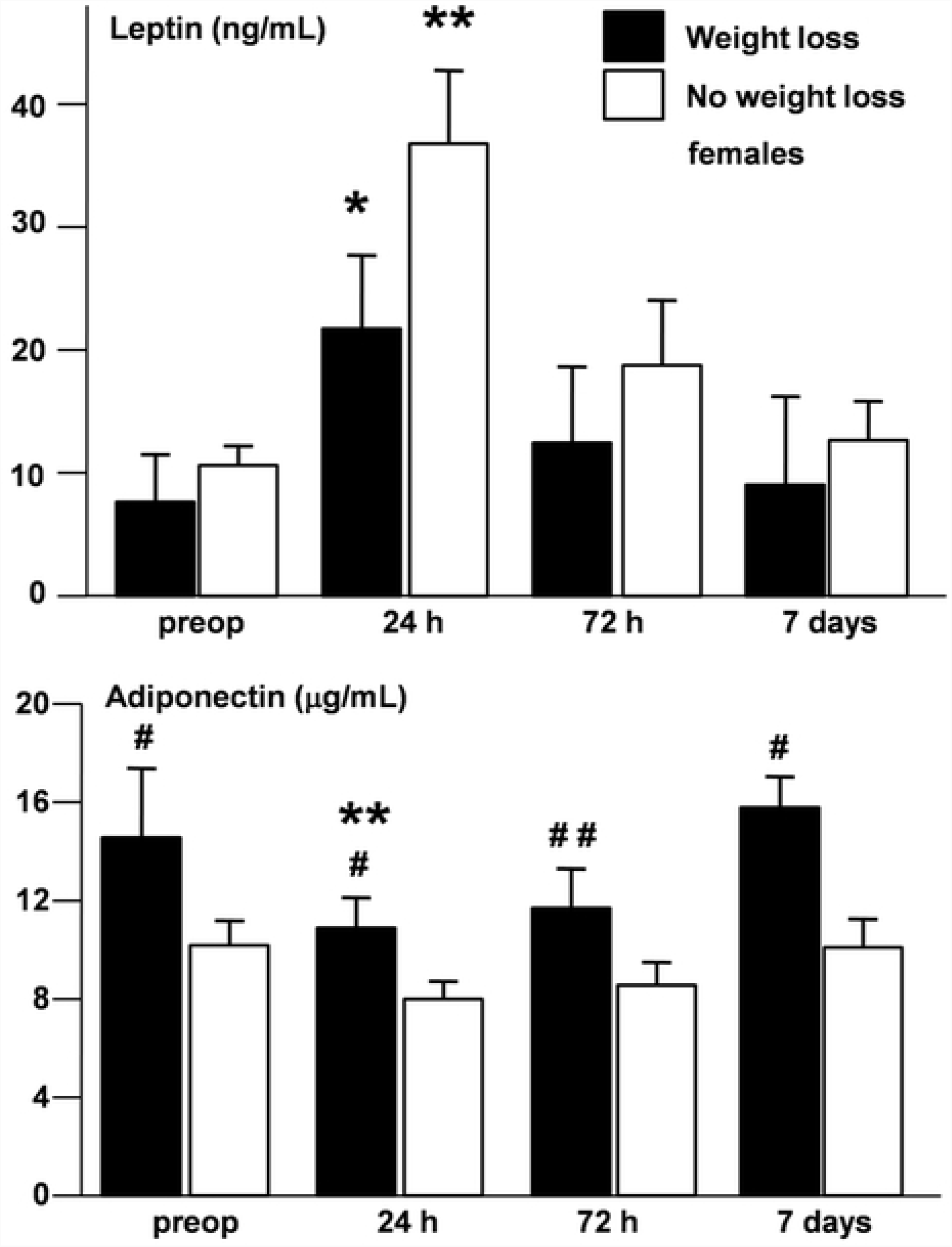
Leptin and adiponectin levels in women with weight loss. Upper panel – leptin, lower panel – adiponectin, white bars – women without weight loss, black bars – women with weight loss, * compared to presurgical levels, # compared to women without weight loss, * or # p < 0.05, ** or ## p < 0.01.

## Discussion

The onset and evolution of CRC are influenced by pleiotropic factors: genetic background [20,25], but also many other variables, such as diet [26], smoking [20] or even fine tuning environment factors such as selenium or other antioxidants [27,28]. The adipocyte is a resource of cytokines and growth stimulating hormones involved in inflammation [1]. Both obesity and an inflammatory environment are correlated with the appearance of various cancers [29-32], being considered risk factors also for CRC [1-3,33]. The majority of our RC patients were indeed overweight or obese (Table 1).

CRC cells possess receptors for leptin and adiponectin [11,34,35]. These adipokines modulate inflammation and display an impact on tumor cell proliferation and angiogenesis as well, both *in vitro* and in animal models. Leptin has oncogenic effects [1,34,37,38], whereas adiponectin has opposite, anti-inflammatory and differentiating effects [1,35,36,38]. Leptin and adiponectin were therefore proposed among vectors linking adiposity and CRC [1-3]. Trying to use serum levels of leptin and adiponectin as prognostic markers in CRC lead, however, to conflicting results. Since leptin proved oncogenic and adiponectin anti-oncogenic effects, one would expect that CRC should be associated with high leptin and low adiponectin levels. Such an adipokinic profile at patients with CRC was indeed described by several authors [1-3,6-8,16,17]. Other studies obtained, however, opposite conclusions, with low leptin [9-13] and/or high adiponectin levels at patients with CRC [15], whereas certain investigators did not observe significant CRC-related changes of leptin [3,14] or adiponectin levels [18,19]. This discordance may be caused by different localizations of CRC in study-enrolled patients [19-22], by cancer staging [8,11,14] and multicenter approach [19-23].

Taking into consideration the high variability of literature data, we wanted to check the modifications of leptin and adiponectin serum concentrations exclusively in rectal cancer. This type of cancer has a greater chance to be diagnosed in pre-metastatic stages [39] and is accompanied by a higher survival rate [40]. We enrolled all our study participants with RC from the same surgical center and compared their leptin and adiponectin levels with weight and age matched healthy controls.

We observed higher leptin and adiponectin serum concentrations in women compared to men in all group pairs. Other authors also described higher levels of these adipokines in women [41,42]. It was therefore important to evaluate adipokine modifications distinctly in men and women, comparing groups of the same gender.

Our RC patients had significantly lower oncogenic leptin and higher anti-oncogenic adiponectin than gender, weight and age matched controls. Such an adipokinic profile was also obtained by certain investigators [9-13,15], in contrast to others [1-3,6-8,14,16-19]. Our results suggest that the adipokinic profile at patients with RC corresponds to an adaptation to this malignant condition rather than an obesity-related triggering factor for malignancy.

We further wanted to investigate the secreting profile of the two adipokines after surgery and observed a significant increase of leptin, as well as a significant decrease of adiponectin 24 hours after surgery in both men and women, followed by a recovery to initial values at 72 hours and 7 days after surgery. Other investigators observed a similar precocious increase of serum leptin and decrease of adiponectin not only after surgery for cancer [43], but also in other acute conditions caused by surgical stress or critical state [44-46]. These modifications are therefore related more with stress by itself and less with the stress triggering factor. The presence and amplitude of these acute changes in adipokine secretion may have a diagnostic or prognostic role (e.g. blunted postoperatory leptin increase meaning the overlap of sepsis and/or a higher mortality risk) [45,46]. Our study could not evaluate the prognostic potential of adipokine measurement, due to the short follow-up period. Although the initial protocol included a serum sample collection also one month after surgery, patients already started dropping out from the study before day 7, forcing its closure. Adipokine concentrations returned nevertheless to presurgical levels before day 7, suggesting that no differences may have been found at a later time point and pleading against an eventual role for these adipokines as parameters for prognostic evaluation.

Preoperatory leptin and adiponectin levels were directly and respectively inversely correlated with BW in both males and females with RC. Fat tissue mass represents an important compartment of the body, which is tightly correlated with and decisively contributes to BW. Leptin is directly secreted by adipose cells and reflects therefore total fat mass in the conditions of stable energy balance [46]. Although also secreted by white adipose cells, adiponectin is inversely correlated to fat mass. The inhibition of adiponectin expression is triggered by higher energy intake rather than by the adiposity per se [47,48].

Various types of stress or inflammatory processes interact with adipokine secretion irrespective of fat mass [43,44,47-49]. We wanted to evaluate therefore the maintenance or distortion of adipokine correlation with body mass by the stress caused by surgical intervention. Surprisingly, whereas the correlation between adipokines and BW was conserved in males, it became non-significant in females not only 24 hours after surgery, but also at later time points, up to one week after surgery. It is not immediately clear why surgical stress causes long-term disappearance of adipokine correlation with BW only in females. Gender differences in adiponectin correlations with BW are probably due to the smaller size of the female subgroup. The correlation slope between leptin and BW was, however, severely modified by surgery in females at all time points when compared to males, where it remained unchanged. The reaction of adipose tissue to surgical stress in RC patients displays therefore a gender divergence. The gender dissimilarities of adipokine association with body mass after stress may be related to differences in hormonal milieu [50] or fat tissue distribution [41,47].

Interestingly, we observed an inverse correlation between leptin and adiponectin before surgery, which became non-significant 24 hours after surgery in both males and females. Significance was restored at 72 hours in females and only after 7 days in males. Since adipose tissue is directly correlated with leptin and inversely correlated with adiponectin [41-43], it is not unexpected to observe an inverse correlation also between leptin and adiponectin. The disturbance of this correlation 24 hours after surgery may be related to the interference of a supplementary factor with adipokine secretion, which is probably stress [43-46].

Leptin, adiponectin and BW were significantly interrelated before surgery in both genders. As described previously, the correlation between BW and adipokines was preserved in males at 24, 72 hours and 7 days after surgery, but became non-significant in females. The three parameters were again all associated significantly in simple correlations at 7 days after surgery only in males. We wanted to observe the interaction of leptin, adiponectin and BW at patients with RC by using hierarchical regression analysis in order to identify the role of BW as a possible confounder between the two adipocytokines according to gender differences.

Hierarchical regression analysis showed the disappearance of significance in the second step (when BW was added) before surgery and after 7 days only in males. In presurgical samples, females preserve the significance of the adiponectin -- leptin association independently of BW (the second step of hierarchical regression). These data point out that the association of leptin with BW is a confounding factor for adiponectin in males, but that adiponectin secretion is influenced by leptin independently of BW in females. Interestingly, although adiponectin and leptin keep being correlated significantly at 72 hours and one week after surgery in females, none of the two adipokines maintain a significant association with BW. These results suggest again that adiponectin and leptin may have a significant interference in women, which is independent of BW. Literature data demonstrate antagonistic effects of leptin and adiponectin [1,3,51,52], but fail to describe a direct interference of any of these adipokines with the secretion of the other [42,53]. Leptin and adiponectin may, however, influence each-other’s levels through opposite modifications in insulin sensitivity [53], although the precise sequence of events is not clear. The direct, BW-independent association of adiponectin with leptin observed only in our group of females, may be explained by the gender variety in insulin sensitivity, probably dependent of gender differences in fat tissue distribution [50,54].

We finally wanted to observe particular adipokine dynamics determined by tumor localization, presurgical nCRT, surgical technique or weight variation and observed significantly higher levels of adiponectin only in women with RC who experienced significant weight loss during the last year before surgery.

Our group was selected from patients who had digestive cancer located exclusively in the rectal region and higher or lower rectal localization did not influence weight variation or adipokinic profile. Other investigators demonstrated a significant impact of chemotherapy on adipokine levels in CRC [4]. The patients included in the respective study had, however, advanced forms of heterogeneously localized CRC and were submitted to chronic, 5 alpha fluorouracil-based palliative chemotherapy. All our patients had only RC and were operated, some being submitted only to a limited period of nCRT before surgery, which did not have a significant impact on adipokine profile. Surgical techniques vary function of tumor localization in RC, with LAR type surgery being accompanied by a less traumatizing and stressful recovery than APR [55]. This well described difference was, however, not reflected by postsurgical adipokine variability, which was similar in patients submitted to LAR or APR. This observation implies that leptin and adiponectin are not suited for discriminating postsurgical evolution of RC patients.

Women who experienced weight loss during one year prior surgery had significantly higher adiponectin levels before surgery and at all time points after surgery. Other authors described leptin and adiponectin modifications with variations in BW [47,48,56-58]. We did not observe differences in leptin levels at women and men who lost weight. Contrary to females, adiponectin levels were not different in men who lost weight. These data plead in favor of gender-determined differences in secreting behavior of adipose tissue with weight loss, as observed by others [57,58]. It is, however, unclear whether this gender difference is related to RC as origin of weight loss or is common to weight variations irrespective of their causes.

Our study has several strengths, but also limitations. Study originality consisted in that all our enrolled patients had a type of digestive cancer localized in the same region – the rectum. The patients were also recruited from the same clinical center and were all operated by the same surgical team, clearly making this study more homogenous than others [2-4,6-9,11,12,17,19-23]. Study drawbacks included the limited number of volunteers, with fewer females with RC and short follow up period due to high dropout rate. This limitation made the evaluation of adipokines as prognostic markers hard to be performed. Evaluated cytokinic spectrum was moreover limited to leptin and adiponectin, other adipokines or fat tissue-related markers (soluble leptin receptor, adiponectin isoforms, resistin, etc.) not being assessed. Finally, fat tissue mass or its distribution was again not assessed at our volunteers.

In conclusion, we observed a particular adipokinic spectrum at RC patients compared to healthy age-, BW- and sex-matched controls, with lower leptin and higher adiponectin, possibly as an adaptation of adipose tissue to this critical situation. Surgical stress produced important modifications of adipokine secretion, which came back to initial values within one week.

Adipokine secretion behaved differently in the two genders suffering of RC. Both leptin and adiponectin levels were higher in women than in men. The correlation between adipokines and BW persisted in men but became non-significant in women up to one week after surgery. Hierarchical regression showed the presence of a significant association between leptin and adiponectin only in females which was independent of BW. The relationship between the two adipokines in females may be mediated by their particular interference with glucose metabolism and insulin sensitivity. Finally, adiponectin was significantly higher in women, but not men with weight loss, when compared to patients without weight loss of the respective gender. Energy metabolism may therefore interfere in another way with adiponectin secretion in females, possibly due to their hormonal background, but also to gender differences in body composition and fat tissue distribution.

These data strengthen the need for further research in evaluating the potential of adipokines as prediction factors in various disease states.

## Acknowledgements.

Part of this work was supported by the European Social Fund through SOP HRD 2007-2013, Priority I “Education and training to support growth and development of knowledge based society” Area of Intervention 1.5 “Doctoral and post-doctoral research support” CERO –CAREER PROFILE: ROMANIAN RESEARCHER, Contract financing: POSDRU/ 159/ 1.5/S/135760.

## References

1. Riondino S, Roselli M, Palmirotta R, Della-Morte D, Ferroni P, Guadagni F. Obesity and colorectal cancer: role of adipokines in tumor initiation and progression. World J Gastroenterol. 2014, 20(18): 5177–5190.

2. Aleksandrova K, Schlesinger S, Fedirko V, Jenab M, Bueno-de-Mesquita B, Freisling H et al. Metabolic mediators of the association between adult weight gain and colorectal cancer: data from the European Prospective Investigation into Cancer and Nutrition (EPIC) cohort. Am J Epidemiol. 2016, 185(9): 751–764.

3. Joshi RK, Lee S-A. Obesity related adipokines and colorectal cancer: a review and meta-analysis. Asian Pac J Cancer Prev. 2014, 15(1): 397–405.

4. Slomian G, Swietochowska E, Nowak G, Pawlas K, Zelazko A, Nowak P. Chemotherapy and plasma adipokines level in patients with colorectal cancer. Postepy Hig Med Dosw. 2017, 71: 281–290.

5. Pietrzyk L, Torres A, Maciejewski R, Torres K. Obesity and obese-related chronic low-grade inflammation in promotion of colorectal cancer development. Asian Pac J Cancer Prev. 2015, 16(10): 4161–4168.

6. Stattin P, Lukanova A, Biessy C, Soderberg S, Palmqvist R, Kaaks R, et al. Obesity and colon cancer: does leptin provide a link? Int J Cancer 2004, 109: 149–152.

7. Ho GYF, Wang T, Gunter MJ, Strickler HD, Cushman M, Kaplan RC, et al. Adipokines linking obesity with colorectal cancer risk in postmenopausal women. Cancer Res. 2012, 72(12): 3029–3037.

8. Guadagni F, Roselli M, Martini F, Spila A, Riondino S, D’Alessandro R, et al. Prognostic significance of serum adipokine levels in colorectal cancer patients. Anticancer Res. 2009, 29: 3321–3328.

9. Arpaci F, Yilmaz MI, Ozet A, Avta H, Ozturk B, Komurcu S, et al. Low serum leptin levels in colon cancer patients without weight loss. Tumori 2002, 88(2): 147–149.

10. Kumor A, Daniel P, Pietruczuk M, Malecka-Panas E. Serum leptin, adiponectin and resistin concentration in colorectal adenoma and carcinoma (CC) patients. Int J Colorectal Dis. 2009, 24(3): 275–281.

11. Erkasap N, Ozkurt M, Erkasap S, Yasar F, Uzuner K, Ihtiyar E, et al. Leptin receptor (Ob-R) mRNA expression and serum leptin concentration in patients with colorectal and metastatic colorectal cancer. Br J Med Biol Res. 2013, 46: 306–310.

12. Hillenbrand A, Fassler J, Huber N, Xu P, Henne-Bruns D, Templin M, et al. Changed adipocytokine concentrations in colorectal tumor patients and morbidly obese patients compared to healthy controls. BMC Cancer 2012, nov. 23 doi: 10.1186/s12885-018-4019-0.

13. Salageanu A, Tucureanu C, Lerescu L, Caras I, Pitica R, Gangura G, et al. Serum levels of adipokines resistin and leptin in patients with colon cancer. J Med Life 2010, 3(4): 416–420.

14. Alexandrova K, Boeing H, Jenab M, Bueno-de-Mesquita HB, Jansen E, van Duijnhoven FJB, et al. Leptin and soluble leptin receptor in risk of colorectal cancer in the European Prospective Investigation into Cancer and Nutrition cohort. Cancer Res. 2012, 72(20): 5328–5337.

15. Tae CH, Kim SE, Jung SA, Joo YH, Shim KN, Jung HK, et al. Involvement of adiponectin in early stage of colorectal carcinogenesis. BMC Cancer 2014 nov 5. doi: 10.1186/1471-2407-14-811.

16. Otake S, Takeda H, Fujishima S, Fukui T, Orii T, Sato T, et al. Decreased levels of plasma adiponectin associated with increased risk of colorectal cancer. World J Gastroenterol. 2010, 16(10): 1252–1257.

17. Touvier M, Fezeu L, Ahluwalia N, Julia C,Charnaux N, Sutton N, et al. Pre-diagnostic levels of adiponectin and soluble vascular cell adhesion molecule-1 are associated with colorectal cancer risk. World J Gastroenterol. 2012, 18(22): 2805–2812.

18. Wei EK, Giovanucci E, Fuchs CS, Willett WC, Mantzoros CS. Low plasma adiponectin levels and risk of colorectal cancer in men: a prospective study. J Natl Cancer Inst. 2005, 97(22): 1688–1694.

19. Nimptsch K, Song M, Aleksandrova K, Katsoulis M, Freisling H, Jenab M, et al. Genetic variation in the ADIPOQ gene, adiponectin concentrations and risk of colorectal cancer – a Mendelian Randomization analysis using data from three large cohort studies. Eur J Epidemiol. 2017, 32(5): 419–430.

20. Liu L, Zhong R, Wei S, Xiang H, Chen J, Xie D, et al. The leptin gene family and colorectal cancer: interaction with smoking behavior and family history of cancer. PLoS One 2013 apr 8. doi:10.1371/journal.pone.0060777

21. Joshi RK, Kim WJ, Lee SA. Association between obesity-related adipokines and colorectal cancer: a case-control study and metaanalysis. World J Gastroenterol. 2014, 20(24): 7941–7949.

22. Aleksandrova K, Drogan D, Boeing H, Jenab M, Bueno-deMesquita HB, Jansen E, et al. Adiposity, mediating biomarkers and risk of colon cancer in the European Prospective investigation into Cancer and Nutrition study. Int J Cancer 2014, 134: 612–621.

23. Song M, Zhang X, Wu K, Ogino S, Fuchs CS, Giovanucci EL, et al. Plasma adiponectin and soluble leptin receptor and risk of colorectal cancer: a prospective study. Cancer Prev Res (Phila) 2013, 6(9): 875–885.

24. Yi L, Ji W, Xiaowei M, Li T, Yanli Y, Chaofan X, et al. A Review of Neoadjuvant Chemoradiotherapy for Locally Advanced Rectal Cancer. Int J Biol Sci. 2016; 12(8): 1022–1031.

25. Brenner H, Kloor M, Pox CP. Colorectal cancer. Lancet. 2014; 383(9927): 1490–1502.

26. Bultman SJ. Interplay between diet, gut microbiota, epigenetic events, and colorectal cancer. Mol Nutr Food Res. 2017 Jan;61(1). doi: 10.1002/mnfr.201500902.

27. Vasiliu I, Preda C, Serban IL, Strungaru SA, Nicoara M, Plavan G, et al. Selenium status in autoimmune thyroiditis. Rev Med Chir Soc Med Nat Iasi. 2015; 119(4): 1037–1144.

28. Preda C, Vasiliu I, Bredetean O, Ciobanu GD, Ungureanu MC, Leustean EL, et al. Selenium in the environment: essential or toxic in human health? Environ Eng Manag J. 2016; 15(4): 913–921.

29. Berger NA. Obesity and cancer pathogenesis. Ann NY Acad Sci. 2014, 1311: 57–76.

30. Azeem S, Gillani SW, Siddiqui A, Jandrajupalli SB, Poh V, Syed Sulaiman SA. Diet and Colorectal Cancer Risk in Asia--a Systematic Review. Asian Pac J Cancer Prev. 2015; 16(13): 5389–5396.

31. Tahergorabi Z, Khazaei M, Moodi M, Chamani E From obesity to cancer: a review on proposed mechanisms. Cell Biochem Funct. 2016; 34(8): 533–545.

32. Diakos CI, Charles KA, McMillan DC, Clarke SJ. Cancer-related inflammation and treatment effectiveness. Lancet Oncol. 2014, 15(11): 493–503.

33. Wang K, Karin M. Tumor-elicited inflammation and colorectal cancer. Adv Cancer Res. 2015, 128: 173–196.

34. Uddin S, Hussain AR, Khan OS, Al-Kuraya KS. Role of dysregulated expression of leptin and leptin receptors in colorectal carcinogenesis. Tumour Biol. 2014; 35(2): 871–879.

35. Byeon JS, Jeong JY, Kim MJ, Lee SM, Nam WH, Myung SJ, et al. Adiponectin and adiponectin receptor in relation to colorectal cancer progression. Int J Cancer. 2010; 127(12): 2758–2767.

36. Kawashima K, Maeda K, Saigo C, Kito Y, Yoshida K, Takeuchi T. Adiponectin and Intelectin-1: important adipokine players in obesity-related colorectal carcinogenesis. Int J Mol Sci. 2017 Apr 19; 18(4). pii: E866. doi: 10.3390/ijms18040866.

37. Zhou W, Tian Y, Gong H, Guo S, Luo C. Oncogenic role and therapeutic target of leptin signaling in colorectal cancer. Expert Opin Ther Targets. 2014; 18(8): 961–971.

38. Adya R, Tan BK, Randeva HS. Differential effects of leptin and adiponectin in endothelial angiogenesis. J Diabetes Res. 2015; 2015:648239. doi: 10.1155/2015/648239.

39. Greco P, Magro G. Pathologic examination and staging of rectal carcinoma: a critical review. Pathologica. 2010; 102(1): 12–27.

40. van der Sijp MP, Bastiaannet E, Mesker WE, van der Geest LG, Breugom AJ, Steup WH, et al. Differences between colon and rectal cancer in complications, short-term survival and recurrences. Int J Colorectal Dis. 2016; 31(10): 1683–1691.

41. Christen T, Trompet S, Noordam R, van Klinken JB, van Dijk KW, Lamb HJ, et al. Sex differences in body fat distribution are related to sex differences in serum leptin and adiponectin. Peptides. 2018; 107:25–31.

42. Ahima RS. Adipose tissue as an endocrine organ. Obesity (Silver Spring). 2006;14 S5: 242S–249S.

43. Teplan V, Senolt L, Hulejova H, Teplan V, Stollova M, Gurlich R. Early changes in serum visfatin after abdominal surgery: a new pro-inflammatory marker in diagnosis? Biomed Pap Med Fac Univ Palacky Olomouc Czech Repub. 2015; 159(3): 489–496.

44. Chachkhiani I, Gürlich R, Maruna P, Frasko R, Lindner J. The postoperative stress response and its reflection in cytokine network and leptin plasma levels. Physiol Res. 2005; 54(3): 279–285.

45. Grigoras I, Branisteanu DD, Ungureanu D, Rusu D, Ristescu I. Early dynamics of leptin plasma level in surgical critically ill patients. a prospective comparative study. Chirurgia (Bucur). 2014; 109(1): 66–72.

46. Florescu A, Bîlha S, Buta C, Vulpoi C, Preda C, Ristescu I, et al. Potential new roles of leptin in health and disease. Rev Med Chir Soc Med Nat Iasi. 2016; 120(2): 252–257.

47. Staiger H, Tschritter O, Machann J, Thamer C, Fritsche A, Maerker E, et al. Relationship of serum adiponectin and leptin concentrations with body fat distribution in humans. Obes Res. 2003; 11(3): 368–372.

48. Musil F, Blaha V, Ticha A, Hyspler R, Haluzik M, Lesna J, et al. Effects of body weight reduction on plasma leptin and adiponectin/leptin ratio in obese patients with type 1 diabetes mellitus. Physiol Res. 2015; 64(2): 221–228.

49. Maslov LN, Naryzhnaya NV, Boshchenko AA, Popov SV, Ivanov VV, Oeltgen PR. Is oxidative stress of adipocytes a cause or a consequence of the metabolic syndrome? J Clin Transl Endocrinol. 2018, 15: 1–5.

50. Selthofer-Relatic K, Radic R, Stupin A, Šišljagić V, Bošnjak I, Bulj N, et al. Leptin/adiponectin ratio in overweight patients - gender differences. Diab Vasc Dis Res. 2018; 15(3): 260–262.

51. Zhang Z, Wang F, Wang BJ, Chu G, Cao Q, Sun BG, et al. Inhibition of leptin-induced vascular extracellular matrix remodelling by adiponectin. J Mol Endocrinol. 2014; 53(2): 145–154.

52. Fenton JI, Birmingham JM, Hursting SD, Hord NG. Adiponectin blocks multiple signaling cascades associated with leptin-induced cell proliferation in Apc Min/+ colon epithelial cells. Int J Cancer. 2008; 122(11): 2437–2445.

53. Yadav A, Kataria MA, Saini V, Yadav A. Role of leptin and adiponectin in insulin resistance. Clin Chim Acta. 2013; 417: 80–84.

54. Bilir BE, Güldiken S, Tunçbilek N, Demir AM, Polat A, Bilir B. The effects of fat distribution and some adipokines on insulin resistance. Endokrynol Pol. 2016; 67(3): 277–282.

55. Smith F, Öhlén J, Persson LO, Carlsson E. Daily Assessment of Stressful events and Coping in early post-operative recovery after colorectal cancer surgery. Eur J Cancer Care (Engl). 2018 Mar;27(2):e12829. doi: 10.1111/ecc.12829.

56. Gajewska J, Ambroszkiewicz J, Klemarczyk W, Chelchowska M, Weker H, Szamotulska K. The effect of weight loss on body composition, serum bone markers, and adipokines in prepubertal obese children after 1-year intervention. Endocr Res. 2018; 43(2):80–89.

57. Ramel A, Arnarson A, Parra D, Kiely M, Bandarra NM, Martinéz JA, et al. Gender difference in the prediction of weight loss by leptin among overweight adults. Ann Nutr Metab. 2010; 56(3):190–197.

58. Reinehr T, Roth C, Menke T, Andler W. Adiponectin before and after weight loss in obese children. J Clin Endocrinol Metab. 2004; 89(8):3790–3794.

